# Soil protist diversity enhances prokaryotic diversity, and regulates dominant prokaryotes and the abundance of key nitrogen cycling genes

**DOI:** 10.1101/2025.02.06.636806

**Authors:** Marta E. Pérez-Villanueva, Stephanie D. Jurburg, Cédric Malandain, Nawras Ghanem, Antonis Chatzinotas

**Affiliations:** Department of Applied Microbial Ecology, Helmholtz Centre of Environmental Research - UFZ, Permoserstrasse 15, 04318, Leipzig, Germany; Hydreka, 51 Av. Rosa Parks, 69009, Lyon, France; German Centre for Integrative Biodiversity Research (iDiv) Halle-Jena-Leipzig, Puschstrasse 4, 04103, Leipzig, Germany; Institute of Biology, Leipzig University, Talstrasse 33, 04103, Leipzig, Germany

**Keywords:** Predator-prey interactions, bacterial community, biodiversity-ecosystem functioning, microbiome, archaea

## Abstract

Soil protists play crucial roles in soil microbial food-webs by preying on bacteria and other microorganisms. However, the effect of protist diversity on soil prokaryotic communities remains poorly understood. This study aimed to elucidate how different protist diversity treatments affect the composition and functionality of soil prokaryotic communities. We established soil microcosms with increasingly complex protist communities, including a control without protists, a medium diversity treatment with three small bacterivorous protists, and a high diversity treatment with seven protists of diverse trophic styles and sizes. Over 21 days, we monitored changes in the prokaryotic community using 16S rRNA gene sequencing and assessed the effects on nitrifiers and denitrifiers by qPCR of nitrogen-cycling genes. Protist diversity explained 23 % of the observed prokaryotic community differentiation over time, with the high-diversity treatment causing the greatest divergence from the control. The most abundant prokaryotes were preferentially predated in all protist treatments. Unexpectedly, the absolute abundance of the *nirK* gene, which is widely distributed among bacterial taxa and thus associated with high functional redundancy, decreased. The differential response of genes with lower distribution and redundancy, such as the bacterial and archaeal *amoA* and the *Nitrospira*-associated *nxrB* genes, to protist diversity indicated selective predation on archaea. High protist diversity systematically enhanced these effects compared to the medium diversity treatment. Overall, protist diversity was positively associated with prokaryotic diversity, which is crucial for maintaining ecosystem stability. These findings highlight the critical role of protist diversity and likely complementary predation in shaping soil prokaryotic communities and their functioning.

## 1. Introduction

Protists are ubiquitous and abundant in soil, with a single gram containing between 10^4^ and 10^8^ protist cells (Adl and Coleman, 2005). Heterotrophic protists consume microorganisms, including bacteria, archaea, fungi, and other protists and play a critical role in soil microbial food-webs (Berger, 1979; Gao et al., 2019; Geisen et al., 2016; Jia et al., 2024; Lee et al., 2022; Lin et al., 2024; Seppey et al., 2017), with direct or indirect effects on plants (Bonkowski, 2004). Predatory protists include various lineages spanning the eukaryotic tree of life (Burki et al., 2020; Keeling and Burki, 2019). Predation mechanisms are shaped by phylogenetically conserved traits such as locomotion type or body size, which influence their food preferences (Flynn et al., 1996; Singh, 1942). Ciliates generate currents and use an expandable oral groove to ingest prey cells, while flagellated protists can achieve high speeds in pursuit of prey, and amoeba use pseudopods to trap and engulf prey (Geisen et al., 2018; Leander, 2020; Nielsen and Kiørboe, 2021). Different protistan feeding patterns can therefore lead to distinct effects on the structure and composition of the soil prokaryotic community, which in turn can trigger changes in its functioning (Amacker et al., 2022; Gao et al., 2019; Nguyen et al., 2023).

Predation is an important cause of bacterial mortality, yet predator-prey interactions contribute to sustain microbial diversity (Burian et al., 2022; Fujino et al., 2023; Gao et al., 2019; Jousset, 2012; Karakoç et al., 2020). Protists are considered generalist predators, because of the large spectrum of species they consume (Geisen et al., 2018; Johnke et al., 2017a, 2014). However, their feeding patterns are often classified as selective because they can discriminate between different prey (Amacker et al., 2020), showing preferences based on prey traits such as size, motility (Glücksman et al., 2010), surface properties (Hoque et al., 2023; Seeger et al., 2010), and even metabolic activity and chemical signals (Jousset et al., 2006; Schulz-Bohm et al., 2017). However, many of these reports are based on single protist experiments and the determinants of predation patterns in communities with higher protist complexity are not yet fully understood.

Alternatively, density-dependent predation contributes to maintaining balance in natural ecosystems, by controlling prey populations and fostering biodiversity (Våge et al., 2018). For example, in marine environments, zooplankton predation on phytoplankton is often density-dependent (Daewel et al., 2014), and similarly, larger terrestrial predators prefer to hunt in areas with higher prey densities (Sinclair et al., 2003). In microbial communities, predation of the protist *Tetrahymena* on the most abundant bacteria has been reported (Saleem et al., 2013), while in soil microcosm experiments with fertilizer application, a decline in the abundance of dominant bacterial phyla was observed in the presence of protists (Asiloglu et al., 2021). Moreover, network analysis suggested a preference of soil protists towards faster growing bacterial prey (Thompson et al., 2021). Although selective predation by protists is well known, we hypothesize that in complex soil communities, predation on more dominant taxa is more relevant for protistan predation patterns, with a positive impact on prokaryotic diversity.

High prokaryotic diversity in soils is closely associated with enhanced soil microbial functionality and functional redundancy, which are critical for ecosystem stability (Allison and Martiny, 2008; Chen et al., 2022b, 2022a; Louca et al., 2018; Maron et al., 2018). Taxonomically distinct microbial taxa within these ecosystems may perform the same functions or metabolic pathways, maintaining ecosystem functioning in the event of species loss and contributing to the buffering capacity against environmental stress (Fetzer et al., 2015; Li et al., 2024). In highly diverse prokaryotic communities, functional redundancy is often promoted through community differentiation driven by factors such as resource partitioning or predator-prey interactions (Louca et al., 2018; Sanders et al., 2018; Wildschutte et al., 2004). However, less redundant genes are more susceptible to extinction when population size declines, for example due to predation, and microbial communities with lower diversity are more likely to have reduced functional redundancy (Louca et al., 2018).

Functional redundancy contributes to maintaining nitrogen cycling in soil (Chen et al., 2022b; Kuypers et al., 2018; Zhu et al., 2024). The nitrogen cycle is highly dependent on prokaryotes, and involves different microbial functional groups that are unevenly distributed within the prokaryotic community (Jiang et al., 2023; Kuypers et al., 2018; Philippot et al., 2007; Prosser and Nicol, 2012).

Nitrification transforms ammonia to nitrite and nitrate through a stepwise process that involves archaeal and bacterial ammonia oxidizers (AOA and AOB respectively) and nitrite oxidizing bacteria (NOB) (Jurburg et al., 2020; Kuypers et al., 2018). These microorganisms are generally slow-growing and highly sensitive to environmental factors such as pH, temperature, and nutrient availability. However, they are less diverse than denitrifiers (Bassin et al., 2015; Daims et al., 2016; Wang et al., 2017), which stepwise convert nitrate to dinitrogen gas. Many bacteria carrying either *nirK* or *nirS* genes are involved in the reduction of nitrite, a central step in denitrification, and are widely distributed in soils (Philippot et al., 2007; Throbäck et al., 2004). Nitrogen cycling genes can therefore serve as a model to understand the effects of predation on microbial functions with different degrees of redundancy (Jurburg and Salles, 2015).

Despite their ecological significance, research on soil protist diversity and its impact on prokaryotic communities remains limited. Due to the high complexity of soil systems, most extant studies use homogeneous laboratory systems (i.e., culture media) (Canter et al., 2018; Carrara et al., 2015; Flues et al., 2017; Friman et al., 2016; Karakoç et al., 2020, 2018; Krumins et al., 2006; Saleem et al., 2013), or rarely consider the effect of more diverse protistan communities (Asiloglu et al., 2021, 2020; Berlinches de Gea et al., 2023; Fujino et al., 2023; Kuppardt et al., 2018). Furthermore, the effect of soil protists on nitrogen cycling functional groups is even less studied (Lin et al., 2024; Pogue and Gilbride, 2007; Wang et al., 2025; Yin et al., 2024).

We developed a soil microcosm experiment with a natural bacterial community exposed to predation pressure with an increasing number and diversity of protist species: a control without protists, a treatment with three bacterivorous flagellate protists and a treatment with seven protists of different trophic modes and sizes. We monitored the prokaryotic community over time using amplicon sequencing of the 16S rRNA gene and its functional profile by quantitative PCR of genes associated with nitrification and denitrification. We hypothesize that protist diversity positively influences soil prokaryotic diversity, with each level of protist diversity shaping a distinct prokaryotic community due to varying predation patterns. Furthermore, we propose that the effects of protist predation on the abundance of prokaryotic functional genes will vary according to their functional redundancy within the microbial community. Specifically, we expect less redundant functions within the nitrogen cycle to be more susceptible to a decline in abundance under increased predation pressure.

## 2. Materials and Methods

### 2.1. Soil microcosms

We used agricultural topsoil (0-20 cm, loamy sand, pH 5.8) obtained in January 2022 from INOQ, GmbH in Schnega, Germany (52°54′26.8″ N, 10°49′20.4″ E), dried it at room temperature, homogenized it with a 2 mm sieve, and stored it at 4 °C. We prepared a prokaryotic soil extract by stirring 300 g of sieved soil in 0.9 % saline solution at a ratio of 1:3.75. We removed larger particles by decanting and then filtering them through a 3 µm nitrocellulose filter. We then removed larger cells such as protists with a second tandem filtration through a 1.2 µm nitrocellulose filter.

We prepared 45 microcosms consisting of 50 g of soil in 100 mL sterile Schott bottles that were first sterilized with γ-radiation (28.79-33.98 kGy, Synergy Health GmbH, Radeberg) prior to inoculation. Then, we inoculated all microcosms with 5.78 mL of the prokaryotic soil extract to achieve 50 % WHC. We homogenized the microcosms, incubated them at 20 °C in the dark for 21 days to allow the prokaryotes to re-colonize the soil prior to protist inoculation, and maintained the water content according to weekly weight determinations throughout the incubation period.

We cultured and maintained in the laboratory seven species of free-living soil protists with various trophic styles, motility modes, and sizes: *Sandona pentamutants* (Sa), *Bodo saltans* (B), *Spumella elongata* (S)*, Cercomonas* sp. (C), *Tetrahymena pyriformis* (T), *Amoeba* sp. (A) and *Rhogostoma pseudocylindrica* (R) (Table 1). *Sandona pentamutants* and *Rhogostoma pseudocylindrica* were obtained from the protist collection of the Institute of Zoology, University of Cologne (Cologne, Germany), provided by Dr. Dumack, and the rest were part of the protist collection of the Department of Applied Microbial Ecology, Helmholtz Centre for Environmental Research - UFZ (Leipzig, Germany). We prepared non-axenic cultures of each protist in inorganic nutrient soytone yeast extract (NSY) (3 g L^-1^) medium (Hahn and Höfle, 1998) at 20-25 °C in the dark and under static conditions, except for *Tetrahymena pyriformis* which was cultured axenically in proteose peptone yeast extract medium (1% proteose peptone, 0.15% yeast extract, 0.01 mM FeCl_3_) (Karakoç et al., 2017). Prior to the experiment and to remove nonspecific debris and smaller cells,

**Table 1.** Traits of the protists added to each protist diversity treatment.

| Protist<sup>1</sup> | Diversity treatment | Trophic mode | Motility mode | Phylum, Class, Family | Size (µm) |
| --- | --- | --- | --- | --- | --- |
| *Sandona pentamutans* (Sa) | Medium, High | Bacterivore | Amoeboflagellar | Cercozoa, Sarcomonadea, *Sandonidae* | ~4-5 |
| *Bodo saltans* (B) | Medium, High | Bacterivore | Flagellar | Euglenozoa, Kinetoplastea, *Bodonidae* | ~4-5 |
| *Spumella elongata* (S) | Medium, High | Bacterivore | Flagellar | Gyrista, Chrysophyceae, *Ochromonadaceaea* | ~6-7 |
| *Cercomonas* sp. (C) | High | Bacterivore | Amoeboflagellar | Cercozoa, Sarcomonadea, *Cercomonadidae* | ~10 |
| *Tetrahymena pyriformis* (T) | High | Bacterivore | Ciliary | Ciliophora, Oligohymenophorea, *Tetrahymenidae* | ~35 |
| *Amoeba* sp. (A) | High | Bacterivore | Amoeboid | Tubulinea, Elardia, *Amoebidae* | ~25 |
| *Rhogostoma pseudocylindrica* (R) | High | Omnivore | Amoeboid | Cercozoa, Thecofilosea, *Rhogostomidae* | ~10-12 |

<sup>1</sup> The abbreviation used for each protist is shown in parenthesis

we concentrated each protist culture by gravity filtration through a 3 μm sterile filter, and resuspended it in 12.6 mL of fresh and filtered (0.2 μm) soil extract.

At the start of the experiment (day 0), we prepared two mixtures of protists to establish the medium and high diversity treatments. The overall volume and total cell number (1x10^4^ cells/g of soil) of the inoculated protists were kept equal in each treatment. The medium protist diversity treatment included the small flagellates Sa, B, and S; the high protist diversity treatment included Sa, B, S, C, T, A, and R and was more diverse in trophic style, motility mode and size than the medium diversity treatment (Table 1). We gently homogenized each mixture by hand shaking and inoculated 1.35 mL of mixture into 15 microcosms per treatment. The same volume of filtered (0.2 μm) soil extract was added to 15 control microcosms without inoculated protists. We adjusted the water content to 65 % WHC with sterile water and incubated the microcosms at 20 °C in the dark. We took destructive samples in quintuplicate at 0, 7 and 21 days after protist inoculation.

### 2.2. Molecular and bioinformatic analyses

We extracted DNA from 0.25 g of each microcosm soil using the ZymoBIOMICS DNA/RNA Miniprep kit (Zymo Research Europe GmbH, Freiburg, Germany) following the manufacturer’s instructions.

We checked the quality and concentration of DNA by electrophoresis on a 1.5 % agarose gel, and with the Qubit™ dsDNA HS Assay Kit (Invitrogen, Waltham, MA, USA), respectively. DNA was stored at −20°C until further use.

We assessed the prokaryotic community in all samples by amplicon sequencing of the 16S rRNA genes. PCR amplifications were performed using a C1000 Touch^TM^ thermal cycler (BIO-RAD, Feldkirchen, Germany), and the amplification was done with 25 cycles of PCR using the optimized primers of the Earth Microbiome Project targeting the V4 hypervariable region of the 16S rRNA gene; 16S_Illu_515F (5’-TCGTCGGCAGCGTCAGATGTGTATAAGAGACAGGTGYCAGCMGCCGCGGTAA-3’) and 16S_Illu_806R (5’-GTCTCGTGGGCTCGGAGATGTGTATAAGAGACAGGGACTACNVGGGTWTCTAAT-3’) (Caporaso et al., 2023). We checked the PCR products by gel electrophoresis. Sequencing was performed on an Illumina MiSeq sequencer (Illumina Inc., San Diego, CA, United States) using the MiSeq Reagent Kit v3 (600 bp).

Raw reads were deposited in the NCBI archive under accession number PRJNA1154586. We conducted the data processing in R v4.4.0 (R Core Team, 2023); we filtered, trimmed, dereplicated, chimera-checked, and merged sequences using DADA2 (v1.32.0) (Callahan et al., 2016) applying the following parameters: *TruncLen =* 245, 205*; maxEE =* 2, 2; *trimLeft* = 10. Taxonomy was assigned to the processed sequences using the SILVA classifier v.138.1 (Quast et al., 2013). The 16S rRNA gene sequenced samples had a range of 28,457-148,722 reads per sample and were rarefied to 28,457 reads per sample (function *rarefy_even_depth*, with seed =1).

#### 2.2.1. Quantitative PCR

We quantified the abundance of the 16S rRNA gene, archaeal and bacterial *amoA* genes, *Nitrospira*-associated *nxrB* gene, and *nirK* gene using a Rotor-Gene 6000 real-time PCR system (QIAGEN GmbH, Hilden, Germany). Primers used, reaction mixtures, and thermocycling conditions for each gene quantification are provided in the Supplementary Material (Table S1). qPCR amplification efficiencies ranged from 82 to 114 %, with R^2^ values of ≥ 0.95. Standard curves were obtained by performing serial dilutions of plasmid vectors that contained amplicons of near full-length target genes.

### 2.3. Data analysis

We processed the sequences and performed the statistical analyses using the R packages *phyloseq* (v.1.48.0) (McMurdie and Holmes, 2013), *vegan* (v.2.6-6.1) (Oksanen et al., 2024), *stats* (v4.4.0) (R Core Team, 2023), *metagMisc* (v.0.5.0) (Mikryukov, 2024), *FSA* (v.0.9.5) (Ogle et al., 2023), *DHARMa* (v.0.4.6) (Hartig, 2022), and *Maaslin2* (v.1.18.0) (Mallick et al., 2021). To assess the differentiation of the prokaryotic community in each protist diversity treatment over time, we performed permutational multivariate analysis of variances (PERMANOVA) using the *adonis2* function from the package *vegan*, followed by pairwise comparisons between protist diversity treatments with the *adonis_pairwise* function from the package *metagMisc*. To account for variation in microbial load across samples, we normalized the abundance of functional genes to the abundance of the 16S rRNA gene, both expressed as the log10 of their gene copies per gram of dry soil. To determine differences between protist diversity treatments in both 16S rRNA gene abundance and relative functional gene abundance we performed repeated Kruskal-Wallis tests per day followed by Dunn’s post hoc test with a Holm correction. We assessed the relationship between 16S rRNA gene abundance and prokaryotic richness or functional gene abundance using the *glm* function of the *stats* package, and evaluated it with the *DHARMa* package. Shapiro and Bartlett tests were performed to check for normality and homogeneity of variance prior to modeling. To investigate the response of the prokaryotic community at the genus level to the different protist diversity treatments, we performed a differential abundance analysis (DAA) using the function *Maaslin2* from the package *Maaslin2*, with the control as the reference group. We attributed differences in the response of prokaryotes to protist diversity, since the total number of inoculated protist cells was set equal for both treatments. We considered significant decreases and increases in the abundance of specific genera compared to the control as negative and positive responses, respectively. Statistical significance was defined as *p*-values < 0.05 for all tests.

## 3. Results

### 3.1. Prokaryotic community differentiation across protist diversity treatments

PCoA analysis of Bray-Curtis dissimilarity between samples (Fig. 1) showed significant differentiation of the prokaryotic community over time (*p*-value = 0.002) and across protist diversity treatments (*p*-value = 0.001), revealing protist diversity as the main driver of the prokaryotic community differentiation, explaining 23 % of it. Both protist diversity treatments resulted in significant shifts in the prokaryotic community on day 7 (*p*-values: Medium = 0.018; High = 0.019) that persisted until day 21 (*p*-values: Medium = 0.018; High = 0.018). On day 7, the differentiation between control and protist diversity treatments (*p*-values: Medium = 0.018; High = 0.021), and between protist diversity treatments (*p*-value: Medium-High = 0.018) was already significant, and by day 21, these differences became even more pronounced (*p*-values: Control-Medium = 0.018; Control-High = 0.019; Medium-High = 0.019).

**Figure 1.**
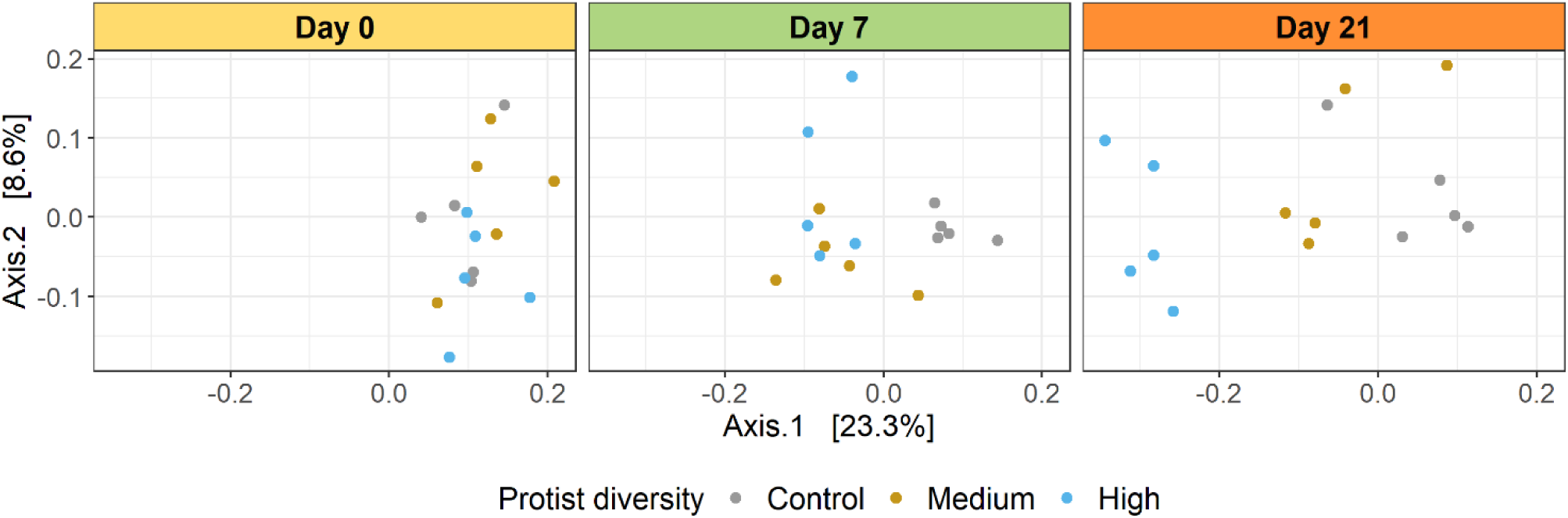
Differentiation of the prokaryotic community over time and across treatments. Principal coordinate analysis (PCoA) plots of Bray-Curtis dissimilarity between samples.

To explore the drivers of this differentiation, we tested for differences in bacterial abundance (16S rRNA gene) between protist diversity treatments, and evaluated the relationship between bacterial abundance and prokaryotic richness using a GLM with a gamma distribution and identity link function. The 16S rRNA gene abundance decreased over time in all treatments (Fig. 2A), with a significant decrease observed in the higher protist diversity treatment compared to the control on days 7 and 21 (*p*-value = 0.017 and *p*-value = 0.022 respectively). In addition, we found a negative relationship between bacterial abundance and richness (β = -193, SE = 34, *p* = 1.52x10^-6^) (Fig. 2B). Pielou’s evenness index showed an increasing trend with increasing protist diversity, but we did not find significant differences between treatments (Fig. S1).

**Figure 2.**
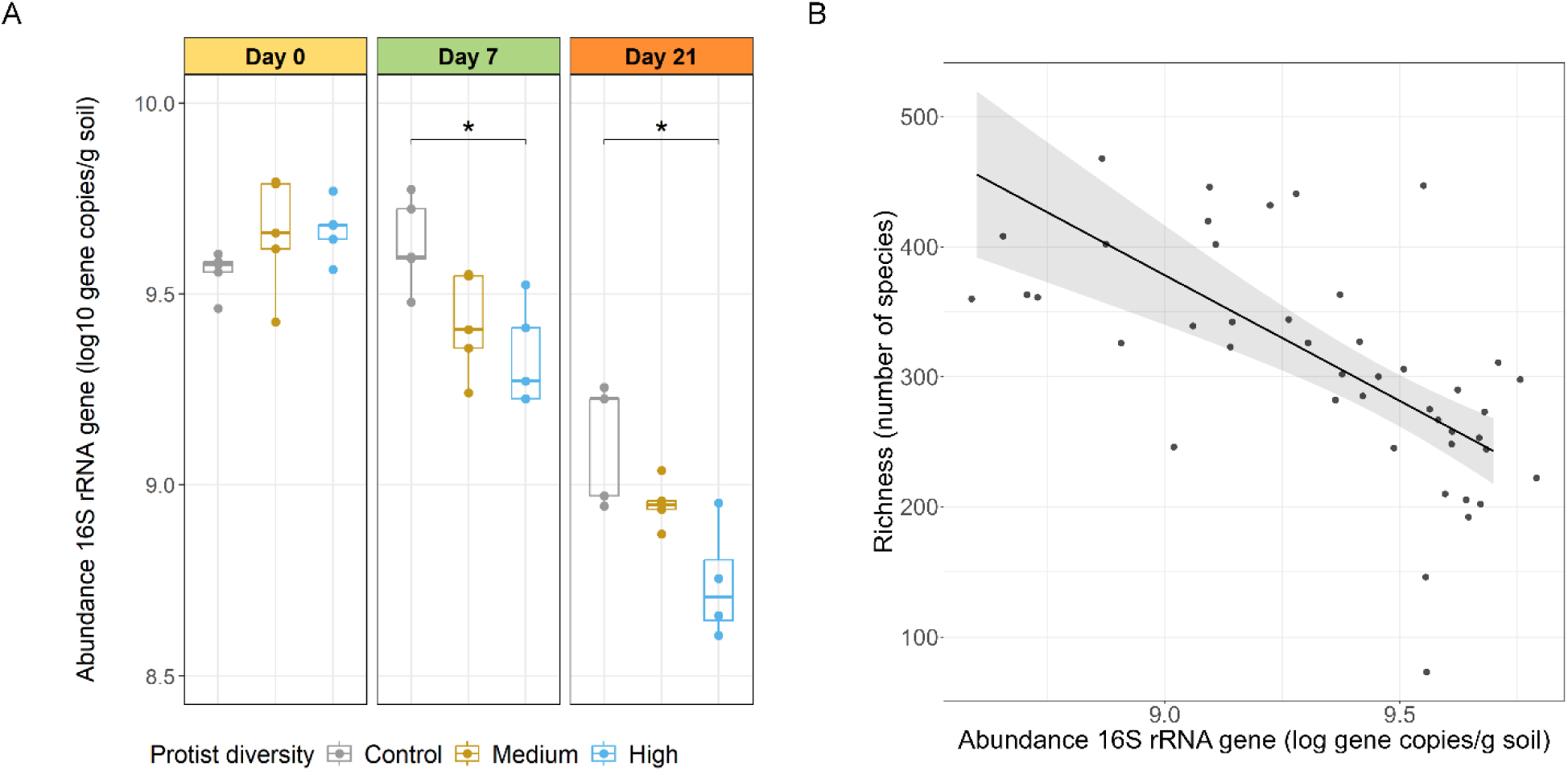
Changes in bacterial abundance and α-diversity over time and across treatments. **A.** Decrease of 16S rRNA gene abundance over time due to protist predation. **B.** Correlation between bacterial abundance and richness. Model estimates for the GLM model correspond to β = -193, SE = 34, *p* = 1.52x10^-6^. For each day, significant differences between treatments were tested separately (Kruskal-Wallis and Dunn’s test) and significant pairwise differences are indicated with brackets (*p*-value < 0.05*).

### 3.2. Differential taxonomic response of soil prokaryotes to protist diversity

In the differential abundance analysis (Fig. 3), we found four genera (0.6 %) that significantly decreased and thirteen genera (1.8 %) that significantly increased in the medium protist diversity treatment relative to the control, corresponding to an average of 6.0 % of the ASVs in the control prokaryotic community. In the high protist diversity treatment, we found three genera (0.4 %) that significantly decreased and 23 (3.2 %) that significantly increased relative to the control, corresponding to 11.6 % of the ASVs in the control prokaryotic community. Only two genera (0.3 %) significantly decreased and five genera (0.7 %) increased in both diversity treatments relative to the control, representing 4.0 % of the ASVs in the control community (Fig. 4A). We found that ASVs affiliated to *Pseudomonas* spp. showed the greatest reduction in relative abundance in the high protist diversity treatment. *Edaphobaculum* spp. and *Mucilaginibacter* spp. responded negatively to both protist diversity treatments. Notably, all negatively affected ASVs belonged to gram-negative phyla and were among the most abundant and prevalent genera in the community (Fig 4B).

**Figure 3.**
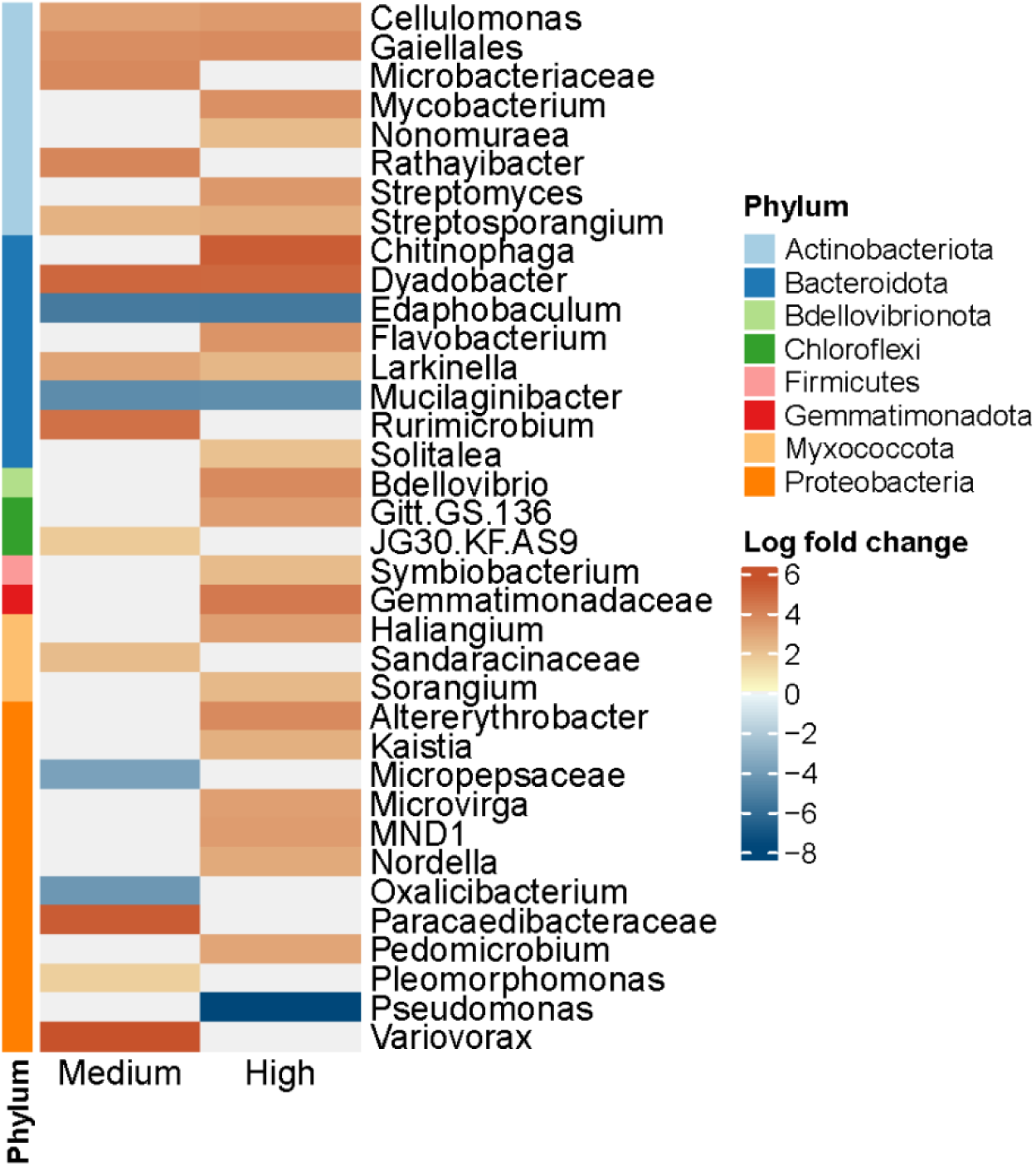
Prokaryotic genera responding to the medium and high protist diversity treatments relatively to the control. The color scale reflects the log fold change in the relative abundance of each genus compared to the control. Shades of red are used for the genera increasing in abundance, shades of blue are used for the genera decreasing in abundance and white color represents no change in the relative abundance. Only genera with significant response for at least one protist diversity treatment are shown (*p*-value < 0.05).

**Figure 4.**
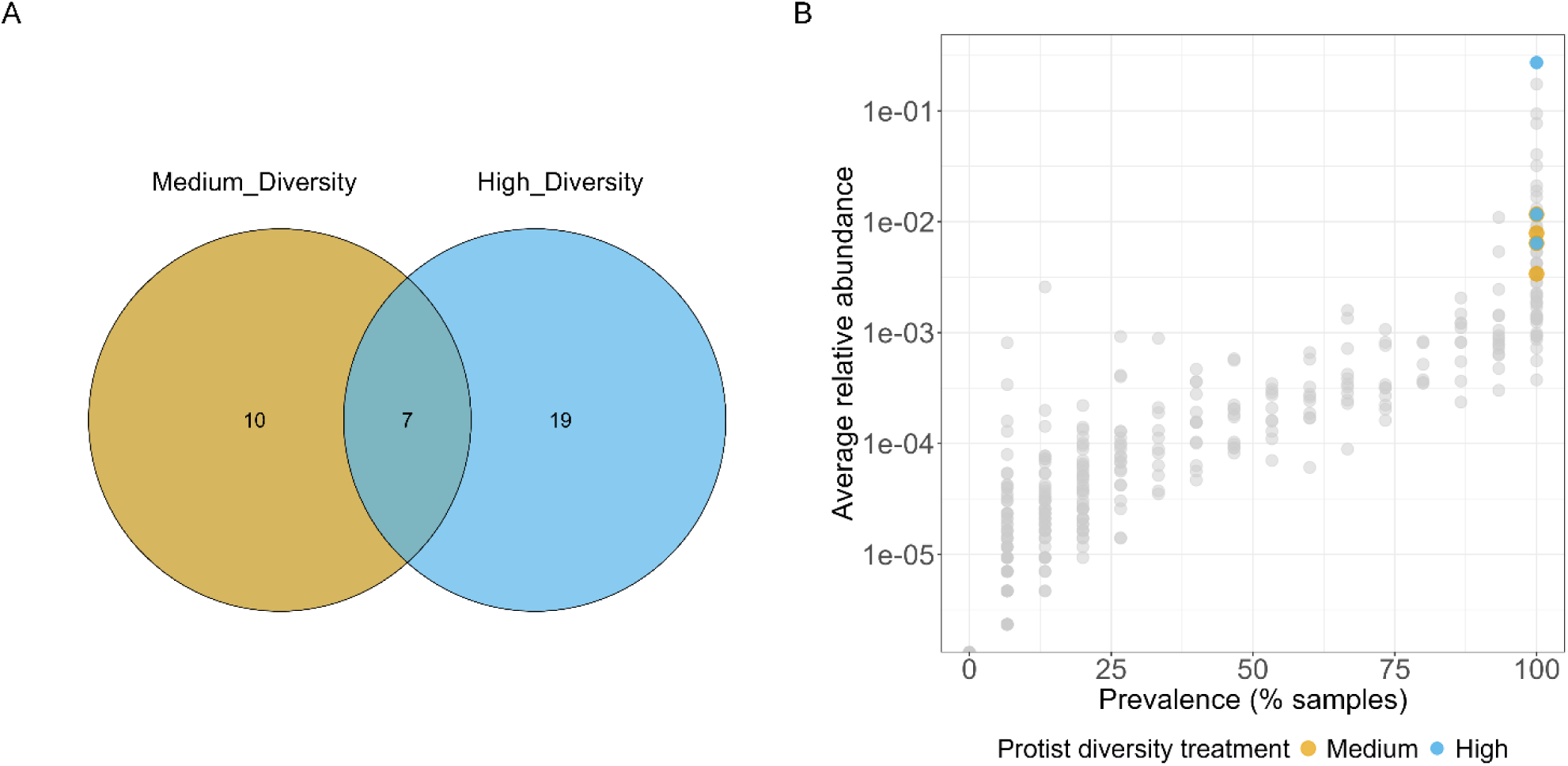
Distribution pattern of prokaryotes significantly affected by the protist diversity treatments. **A.** Venn’s diagram showing the number of prokaryotic genera responding to each protist diversity treatment relative to the control. **B.** Prevalence distribution of genera that responded negatively to protist diversity treatments. Gray dots represent prokaryotic genera in the control, yellow and light-blue dots indicate prokaryotic genera responding negatively to the medium and high protist diversity treatment, respectively.

### 3.3. Effect of protist diversity on the abundance of nitrifier and denitrifier genes

The relative abundances of archaeal *amoA* and *nirK* genes increased significantly on day 7 in the high protist diversity treatment compared to the control (*amoA*: *p*-value = 0.004; *nirK*: *p*-value = 0.03) (Fig. 5). In the case of the *nirK* gene the increase in relative abundance remained significant on day 21 (*p*-value = 0.02). We did not observe significant differences in relative abundances between protist diversity treatments for either the bacterial *amoA* gene or the *Nitrospira*-associated *nxrB* gene. In addition, to determine whether there was a correlation between the changes in bacterial abundance and the abundance of functional genes, we performed a GLM with a gamma distribution and identity link function. We found a positive relationship between the abundance of the 16S rRNA gene and the archaeal *amoA* (β = 0.28, SE = 0.04, *p* = 2.2 x 10^-8^) and the *nirK* (β = 0.34, SE = 0.04, *p* = 4.26 x 10^-11^) genes. In contrast, no significant relationship was found for the bacterial *amoA* and the *Nitrospira*-associated *nxrB* gene (Fig. 5). The *Nitrobacter*-associated *nxrB* gene had a similar response as the *Nitrospira*-associated *nxrB* gene (Fig. S2), however, it was not possible to fit the model for the relationship.

**Figure 5.**
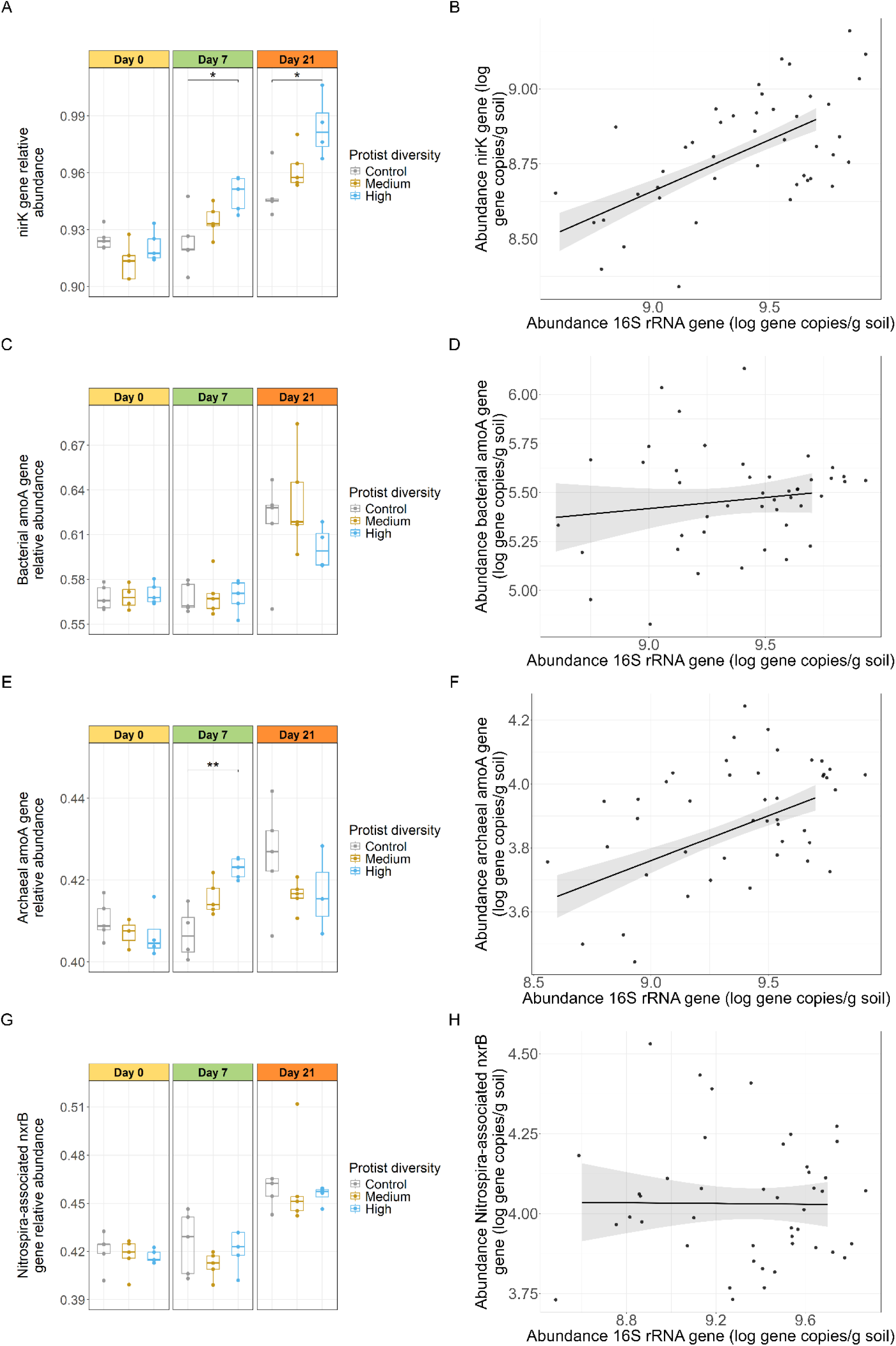
Effect of protist diversity on the abundance of functional genes. **A-B.** *nirK* gene, **C-D.** bacterial *amoA* gene, **E-F.** archaeal *amoA* gene, **G-H.** *Nitrospira*-associated *nxrB* gene. On the left panel is displayed the relative abundance of the functional gene per day and protist diversity treatment, and on the right panel the relationships between absolute abundances of the 16S rRNA gene and the respective functional gene. Model estimates for the GLM models correspond to (B) β = 0.34, SE = 0.04, *p* = 4.3x10^-11^ ***, (D) β = 0.11, SE = 0.11, *p* = 0.29, (F) β = 0.28, SE = 0.04, *p* = 2.2x10^-8^ ***, (H) β = -0.006, SE = 0.07, *p* = 0.94. For each day, significant differences between treatments were tested separately (Kruskal-Wallis and Dunn’s test) and significant pairwise differences are indicated with brackets (*p*-value < 0.05*, 0.01**).

## 4. Discussion

### 4.1. Protist diversity shapes soil prokaryotic communities through predation

Our results are consistent with reports on the role of protists in shaping experimental communities through predation (Amacker et al., 2022; Flues et al., 2017; Fujino et al., 2023; Gao et al., 2019; Glücksman et al., 2010; Hoque et al., 2023). We demonstrated that protist diversity was the main driver of the prokaryotic community diversification, which diverged most strongly in the high protist diversity treatment. As protist diversity increased, 16S rRNA gene abundance decreased and prokaryotic richness increased. This is in line with studies using simplified microcosm communities, which have shown that predation by multiple protists can lead to increased prey richness (Corno et al., 2008; Saleem et al., 2012), one explanation being increasing complementarity effects and better resource use (Filip et al., 2014; Moorthi et al., 2008; Saleem et al., 2012).

The biodiversity-ecosystem function (BEF) theory posits that species diversity improves resource use through complementarity or selection effects (Cardinale et al., 2002). Complementarity effects arise from processes such as niche differentiation, resource partitioning, and positive interactions between species (Brooker et al., 2021; Eisenhauer, 2012; Loreau and Hector, 2001; Saleem et al., 2012). As selective feeding on bacterial prey by protists can result in species-specific effects on the prey (Amacker et al., 2022; Bell et al., 2010; Glücksman et al., 2010; Saleem et al., 2012), complementarity effects due to differences in feeding modes and efficiencies are not unexpected in diverse protist communities. Although we did not directly measure complementarity or selection effects, the observed increase in prokaryotic richness with higher protist diversity, combined with the different trophic modes, motility strategies and sizes of protists in the high diversity treatment, supports the idea of niche differentiation and different resource access. This suggests that complementarity effects were likely involved in shaping community dynamics in our microcosms.

However, such diversity effects via feeding complementarity may not be relevant in high diversity systems which go beyond the protist diversity established in our microcosms (Berlinches de Gea et al., 2023).

We also found that a higher number of prokaryotes responded positively to the protist diversity compared to those that responded negatively. This response was more pronounced in the high protist diversity treatment and is consistent with our findings on prokaryotic richness. In addition to complementary and selection effects associated to protist diversity, consumption of microorganisms by protists leads to the recycling and the release of nutrients (Adl and Gupta, 2006; Gao et al., 2019), which can potentially promote growth of other bacteria and plants (Bonkowski, 2004; Clarholm, 1985; Xiong et al., 2020).

Interestingly, the bacteria that became less abundant in both protist diversity treatments were gram negative bacteria. Members of this group are more likely to be a suitable prey for protists than gram positive bacteria, probably due to the more complex cell-wall structure of the latter (Chandarana and Amaresan, 2022; Murase et al., 2006). Among the significantly affected bacterial taxa, non-motile bacteria (*Edaphobaculum* spp. and *Mucilaginibacter* spp.) were similarly predated in both protist diversity treatments, whereas *Micropepsaceae* spp., *Oxalicibacterium* spp., and *Pseudomonas* spp. were affected in either medium or high protist diversity treatments. Susceptibility to predation by protists may be influenced by predator competition for food (Jousset, 2012; Saleem et al., 2013), as well as anti-predation mechanisms of the prey, with variable efficacy against different predators (Jousset, 2012). Enhanced motility is an important mechanism in bacteria to escape predation (Amacker et al., 2020; Matz and Jürgens, 2003). However, the development of additional anti-predation strategies is often required to further enhance their resistance to predation. For example, motile bacteria from the genus *Pseudomonas* have shown variable susceptibility to predation, which can be enhanced by, for example, the production of exometabolites (Amacker et al., 2020; Jousset et al., 2006) or cooperative predation defense (Zhang et al., 2021).

Genera that responded negatively to the protist treatments belonged to the most abundant and prevalent genera in the control community, suggesting a density-dependent negative response to protist diversity, and a predation pattern towards dominant prokaryotes. Saleem et al. (2013) previously observed preferential predation of *Tetrahymena* sp. on more dominant bacterial species in a simplified microcosm experiment. They also reported that *Tetrahymena* sp. can dominate the predator community due to high growth rate. Even though this protist was present in our high protist diversity treatment, our soil experiment also showed a density-dependent predation in the intermediate diversity that did not include *Tetrahymena*, suggesting that density-dependent predation is a common strategy in complex soil protistan communities. This predation pattern of protists has previously been observed in experiments simulating riverine environments or waterlogged soils (Batani et al., 2016; Murase et al., 2006). Predation on dominant taxa creates niches for less competitive, rare taxa to thrive, reducing competitive exclusion, contributing to an increase in richness and likely promoting evenness within the community (Jousset et al., 2017; Kurm et al., 2019; Rodriguez-Valera et al., 2009; Saleem et al., 2012). Interestingly, we found an increase in the relative abundance of *Bdellovibrio* sp. in the high protist diversity compared to the control, but not at the medium diversity. *Bdellovibrio* is an obligate predator of gram-negative bacteria in a wide range of habitats (Bratanis et al., 2020; Johnke et al., 2017b). This suggests that *Bdellovibrio* spp. may benefit from the presence of either specific protists or higher protistan diversity in soil.

### 4.2. Protist diversity influences the abundance of nitrogen cycling genes

Protist predation in soil has been most studied in the context of general community changes and plant health (Asiloglu et al., 2020; Bonkowski, 2004; Fujino et al., 2023; Rosenberg et al., 2009), but its specific effects on the abundance of functional groups remain underexplored (Jiang et al., 2023; Lin et al., 2024; Wang et al., 2025; Yin et al., 2024). We assessed the effect of protist diversity on the abundance of genes involved in key steps of the nitrogen cycle, which are differently distributed in the prokaryotic community. The *nirK* gene, which encodes a key enzyme in denitrification, is abundant in soil, as approximately 5 % of the total soil microbial community are denitrifiers and carry either the *nirK* or *nirS* gene (Demanèche et al., 2009). We observed significant differences in the relative abundance of *nirK* between the control and the high protist diversity treatment, and a positive correlation with the 16S rRNA gene abundance. This finding suggests that the reduction in the relative abundance of the *nirK* gene is a cascading effect driven by the reduction in the overall bacterial abundance, likely due to predation. Although redundant functional genes, such as *nirK*, are often considered less susceptible to changes in community composition due to their widespread distribution (Louca et al., 2018; Philippot et al., 2013), our findings suggest that density-dependent predation may also negatively affect abundant genes in the microbiome, consistent with the observed effects on the overall bacterial abundance. Similarly, Yin et al. (2024) observed a regulation of denitrification by protists that resulted in higher protist growth rates. Interestingly, despite a decrease in absolute abundance, the relative abundance of *nirK* increased over time, especially in the high diversity treatment, suggesting functional resilience that preserves the denitrification potential under predation pressure.

In contrast, nitrification involves more specialized microbial groups, with key genes such as *amoA* (ammonia monooxygenase) and *nxrB* (nitrite oxidoreductase) considered to be less redundant in soil microbial communities (Wang et al., 2022). However, the abundance of bacterial *amoA* and *Nitrospira*-associated *nxrB* genes did not correlate with changes in 16S rRNA gene abundance across protist diversity treatments, likely because the density-dependent predation pattern may provide an indirect protection for bacteria with these genes, which are less represented in the microbiome. However, archaeal *amoA* gene abundances were positively correlated with 16S rRNA gene abundance and showed significant differences between treatments at day 7. This suggests distinct dynamics of protist interactions with ammonia oxidizing archaea (AOA) and bacteria (AOB).

Similar to *nirK*, we found significant differences in the relative abundance of archaeal *amoA* genes between protist diversity treatments, but bacterial *amoA* remained unaffected, suggesting selective predation. Predation preferences can be influenced by traits such as cell size, cell composition, motility and microbial volatiles, as well as environmental factors like soil aggregate structure, oxygen availability, and nutrient gradients (Ballen-Segura et al., 2016; Erktan et al., 2020; Roberts et al., 2011; Schulz-Bohm et al., 2017). Moreover, recent studies indicate that protists interact differently with AOA and AOB, with a higher frequency of negative interactions with AOA, regardless of additional treatments such as fertilization regimes (Jia et al., 2024; Lin et al., 2024). These interactions appear to be identity dependent; for example, amoebae from the Variosea group dominated the interactions with ammonia-oxidizing microorganisms in an acidic soil (Lin et al., 2024), while Cercozoa emerged as the protist phylum that consistently suppressed AOA in a fertilization experiment where other protists phyla were also identified as dominant (Jia et al., 2024). In addition, evidence from aquatic and grassland ecosystems suggest preferential protist predation on archaeal taxa, although the underlying factors remain unclear (Ballen-Segura et al., 2016; Gu et al., 2023).

Significant differences in archaeal *amoA* gene abundance were only observed in the high protist diversity treatment, potentially indicating enhanced interactions between AOA and the protists in this treatment, and supporting the ideas of competition at the predator level and resource partitioning under high protist diversity (Johnke et al., 2017a). Concurrently, at the prey level, higher predation pressure may increase competition between prokaryotic taxa. This, combined with an increased nutrient availability due to predation, may negatively affect the abundance of AOA, which are favored in nutrient-poor environments (Amacker et al., 2020; Batani et al., 2016; Prosser and Nicol, 2012). These results suggest that protist predation on soil prokaryotic communities predominantly follows a density-dependent pattern, with more abundant taxa being targeted. Contrary to our hypothesis, functional genes with lower redundancy appear to be less susceptible to the negative effects of protist predation, particularly in soils with diverse protist communities. However, evidence for selective predation on ammonia-oxidizing archaea was also observed, albeit to a lesser extent.

## 5. Conclusions

Our study extends the understanding of how different predation pressure by protists affects soil prokaryotic communities and their functionality. Our results show that, although protists may have different feeding patterns, the most abundant prokaryotes were consistently the preferred target of predation. Surprisingly, this predation negatively affected the absolute abundance of genes with high functional redundancy and distribution among microbial communities, but did not significantly affect low-redundant genes, which are considered more susceptible to extinction. However, the relative abundances of nitrogen-cycling genes indicated that protist predation did not compromise the overall functional potential. Moreover, our results suggest selective protist predation on ammonia-oxidizing archaea. Overall, high protist diversity had the strongest effect on promoting prokaryotic diversity and on the functionality, highlighting its ecological importance in shaping soil microbial ecosystems. Future research should further investigate how environmental stressors affect protist diversity and subsequently cascade down to the prokaryotic community, as protists may be more sensitive to changes in the environment than fungal and bacterial communities (Degrune et al., 2024; Zhao et al., 2019).

## CRediT authorship contribution statement

**Marta E. Pérez-Villanueva:** Writing – original draft, Writing – review & editing, Conceptualization, Investigation, Data curation, Visualization, Software, Methodology, Formal analysis. **Stephanie D. Jurburg:** Writing – review & editing, Methodology, Conceptualization, Supervision. **Cédric Malandain:** Writing – review & editing, Supervision, Conceptualization, Funding acquisition. **Nawras Ghanem:** Writing – review & editing, Supervision, Conceptualization. **Antonis Chatzinotas:** Writing – review & editing, Supervision, Funding acquisition, Conceptualization, Methodology.

## Data availability

16S rRNA gene sequencing raw data were deposited into NCBI Sequence Read Archive (SRA) under the accession number PRJNA1154586. qPCR raw data were deposited Zenodo repository (https://doi.org/10.5281/zenodo.14647026)

## Acknowledgements

We thank Nicole Steinbach and Anett Heidtmann for their excellent technical assistance, Dr. Ulrike Schlägel for statistical advice (all UFZ), PD Dr. Kenneth Dumack from the University of Cologne for providing two protist strains and Prof. Graeme Nicol and Dr. Christina Hazard from École Central de Lyon for providing the plasmid vectors used in the standard curves for some of our qPCR quantifications. This work was supported by the EU’s H2020 research and innovation program under the Marie Skłodowska-Curie grant agreement No. 956496 MSCA-ITN-H2020 project ARISTO.

