## Supplementary material for "Soil protist diversity enhances prokaryotic diversity, and regulates dominant prokaryotes and the abundance of key nitrogen cycling genes"

Table S1. Primers, primers sequences, thermal conditions and reaction mixtures used for qPCR

| **Target gene** | **Primer name** | **Primers sequence** | **Thermal cycling conditions** | **Reaction mixture^1,2,3^** | **Reference** |
| --- | --- | --- | --- | --- | --- |
| 16S rRNA | 926F | 5’-AAACTCAAAKGAATTGACGG-3’ | 95 °C for 3’ and 45 cycles of 95 °C for 5”, 61 °C for 10”, 72 °C for 10” | 10 μl of Master Mix (2X), 1.2 μl of each primer, 2 μl of template DNA and 5.6 μl of sterile H_2_O | Bacchetti De Gregoris et al., 2011 |
|  | 1062R | 5’-CTCACRRCACGAGCTGAC-3’ |  |  |  |
| Bacterial *amoA* | amoA1F | 5’-GGGGTTTCTACTGGTGGT-3’ | 95 °C for 15’, 35 cycles of 94 °C for 15”, 55 °C for 30”, 72 °C for 30”, and 82 °C for 5” | 10 μl of Master Mix (2X), 2.4 μl of each primer, 2 μl of template DNA and 3.2 μl of sterile H_2_O | Rotthauwe et al., 1997 |
|  | amoA2R | 5’-CCCCTCKGSAAAGCCTTCTTC-3’ |  |  |  |
| Archaeal *amoA* | crenamoA23F | 5’-ATGGTCTGGCTWAGACG-3’ | 95 °C for 15’ and 35 cycles of 94 °C for 15”, 56 °C for 30”, 72 °C for 30” | 10 μl of Master Mix (2X), 1.2 μl of each primer, 2 μl of template DNA and 5.6 μl of sterile H_2_O | Tourna et al., 2008 |
|  | crenamoA616R | 5’-GCCATCCATCTGTATGTCCA-3’ |  |  |  |
| Nitrospira’s *nxrB* | nxrB169F | 5’-TACATGTGGTGGAACA-3’ | 95 °C for 5’, 35 cycles of 95 °C for 40”, 56 °C for 40”, 72 °C for 90”, and 72 °C for 10’ | 10 μl of Master Mix (2X), 0,6 μl of each primer, 2 μl of template DNA and 6.8 μl of sterile H_2_O | Pester et al., 2014 |
|  | nxrB638R | 5’-CGGTTCTGGTCRATCA-3’ |  |  |  |
| Nitrobacter’s *nxrB* | nxrB-1F | 5’-ACGTGGAGACCAAGCCGGG-3’ | 95 °C for 5’, 35 cycles of 95 °C for 60”, 55 °C for 60”, 72 °C for 120”, and 72 °C for 10’ | 10 μl of Master Mix (2X), 0.6 μl of each primer, 2 μl of template DNA and 6.8 μl of sterile H_2_O | Vanparys et al., 2007 |
|  | nxrB-1R | 5’-CCGTGCTGTTGAYCTCGTTGA-3’ |  |  |  |
| *nirK* | nirK-876 | 5’-ATYGGCGGVCAYGGCGA-3’ | 95 °C for 3’ and 45 cycles of 95 °C for 5”, 60 °C for 15”, 72 °C for 10” | 10 μl of Master Mix (2X), 1.2 μl of each primer, 2 μl of template DNA and 5.6 μl of sterile H_2_O | Henry et al., 2004 |
|  | nirK-1040 | 5’-GCCTCGATCAGRTTRTGGTT-3’ |  |  |  |

^1^Master Mix: SensiFAST^TM^ SYBR ® qPCR Master Mix (2X) (MeridianBioscience, London, UK).

^2^Primer solutions were prepared at 10 μM.

^3^Template DNA was prepared at 2.5 ng/uL for all samples.

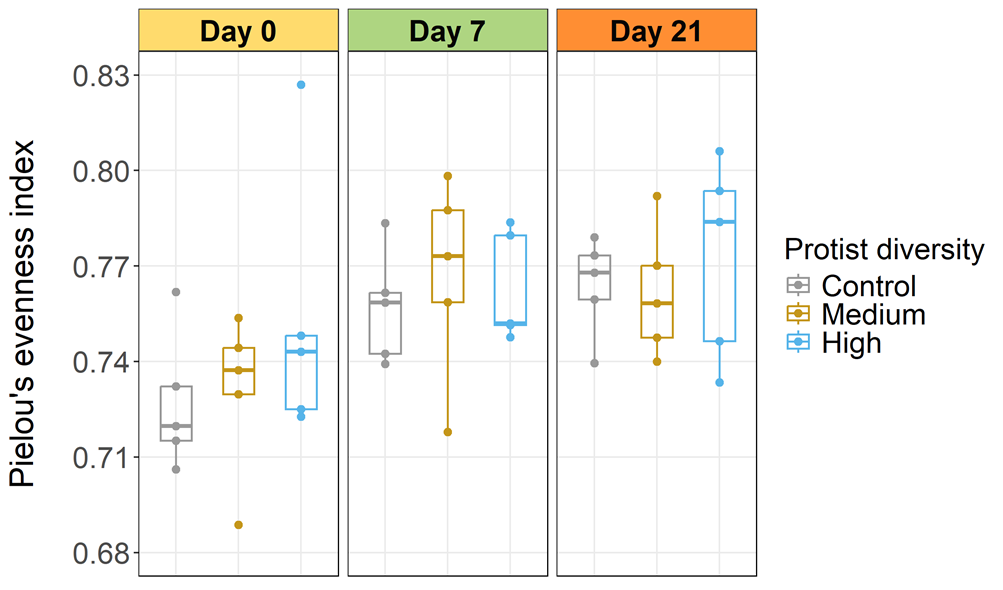

Figure S1. Pielou’s evenness index for each protist diversity treatment over time.

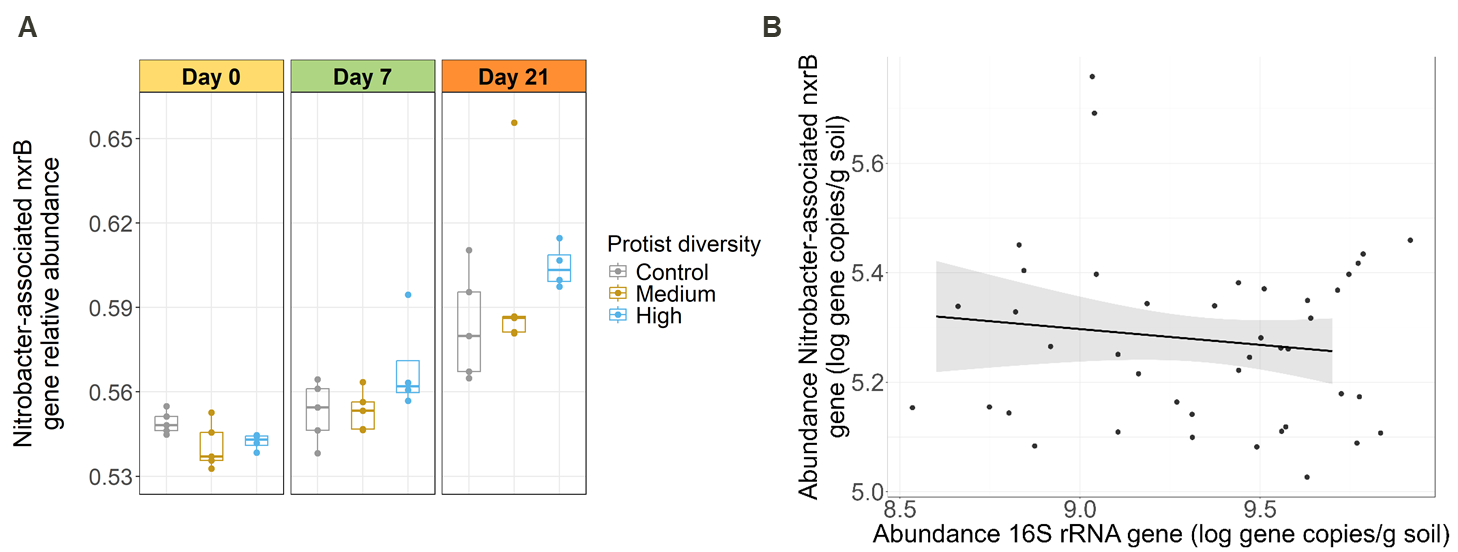

Figure S2. Effect of protist diversity on the abundance of the *Nitrobacter*-associated *nxrB* gene. **A.** *Nitrobacter*-associated *nxrB* gene relative abundance. **B.** Relationship between 16S rRNA gene and *Nitrobacter*-associated *nxrB* gene absolute abundances per day and protist diversity treatment. No significant differences were found between treatments per day (Kruskal-Wallis and Dunn’s test). It was not possible to fit the selected model for the relationship.
